# The Field-Dependent Nature of PageRank Values in Citation Networks

**DOI:** 10.1101/2023.01.05.522943

**Authors:** Benjamin J. Heil, Casey S. Greene

## Abstract

The value of scientific research can be easier to assess at the collective level than at the level of individual contributions. Several journal-level and article-level metrics aim to measure the importance of journals or individual manuscripts. However, many are citation-based and citation practices vary between fields. To account for these differences, scientists have devised normalization schemes to make metrics more comparable across fields. We use PageRank as an example metric and examine the extent to which field-specific citation norms drive estimated importance differences. In doing so, we recapitulate differences in journal and article PageRanks between fields. We also find that manuscripts shared between fields have different PageRanks depending on which field’s citation network the metric is calculated in. We implement a degree-preserving graph shuffling algorithm to generate a null distribution of similar networks and find differences more likely attributed to field-specific preferences than citation norms. Our results suggest that while differences exist between fields’ metric distributions, applying metrics in a field-aware manner rather than using normalized global metrics avoids losing important information about article preferences. They also imply that assigning a single importance value to a manuscript may not be a useful construct, as the importance of each manuscript varies by the reader’s field.

## Introduction

There are more academic papers than any human can read in a lifetime. Attention has been given to ranking papers, journals, or researchers by their “importance,” assessed via various metrics. Citation count assumes the number of citations determines a paper’s importance. The h-index and Journal Impact Factor focus on secondary factors like author or journal track records. Graph-based methods like PageRank or disruption index use the context of the citing papers to evaluate an article’s relevance [1,2,3,4]. Each of these methods has limitations, and permutations exist that attempt to shore up specific weaknesses [5,6,7,8].

One objection to such practices is that “importance” is subjective. The San Francisco Declaration on Research Assessment (DORA) argues against using Journal Impact Factor, or any journal-based metric, to assess individual manuscripts or scientists [9]. DORA further argues in favor of evaluating the scientific content of articles and notes that any metrics used should be article-level (https://sfdora.org/read/). However, even article-level metrics often ignore that the importance of a specific scientific output will fundamentally differ across fields. Even Nobel prize-winning work may be unimportant to a cancer biologist if the prize-winning article is about astrophysics.

Because there are differences between fields’ citation practices [10], scientists have developed strategies including normalizing the number of citations based on nearby papers in a citation network, rescaling fields’ citation data to give more consistent PageRank results, and so on [5,11,12,13]. Such approaches normalize away field-specific effects, which might help to compare one researcher with another in a very different field. However, they do not address the difference in the relevance of a topic between fields. This phenomenon of field-specific importance has been observed at the level of journal metrics. Mason and Singh recently noted that depending on the field, the journal *Christian Higher Education* is either ranked as a Q1 (top quartile) journal or a Q4 (bottom quartile) journal [14].

It is possible that, while global journal-level metrics fail to capture field-specific importance, article-level metrics are sufficiently granular that the importance of a manuscript remains constant across fields. We investigate the extent to which article-level metrics generalize between fields. We examine this using MeSH terms to define fields and use field-specific citation graphs to assess their importance within the field. While it is trivially apparent that journals or articles that do not have cross-field citations will have variable importance, we ignore these cases. We include only those including citations in both fields, where we expect possible consistency. We first replicate previous findings that journal-level metrics can differ substantially among fields. We also find field-specific variability in importance at the article level. We make our results explorable through a web app that shows metrics for overlapping papers between pairs of fields.

Our results show that even article-level metrics can differ substantially among fields. We recommend that metrics used for assessing research outputs include field-specific, in addition to global, ones. While qualitative assessment of the content of manuscripts remains time-consuming, our results suggest that within-field and across-field assessment remains key to assessing the importance of research outputs.

## Results

### Journal rankings differ between fields

In an attempt to quantify the relative importance of journals, scientists have created rankings using metrics the Journal Impact Factor, which essentially uses citations per article, and those that rely on more complex representations like Eigenfactor [15]. Previous reports note that journal rankings differ substantially between fields using metrics based on citation numbers [14]. We calculated a field-specific PageRank-based score for each journal as the median PageRank of manuscripts published in that journal for that field (Fig. 1 A). We first sought to understand the extent to which PageRank replicated journal ranking differences across fields.

**Figure 1:**
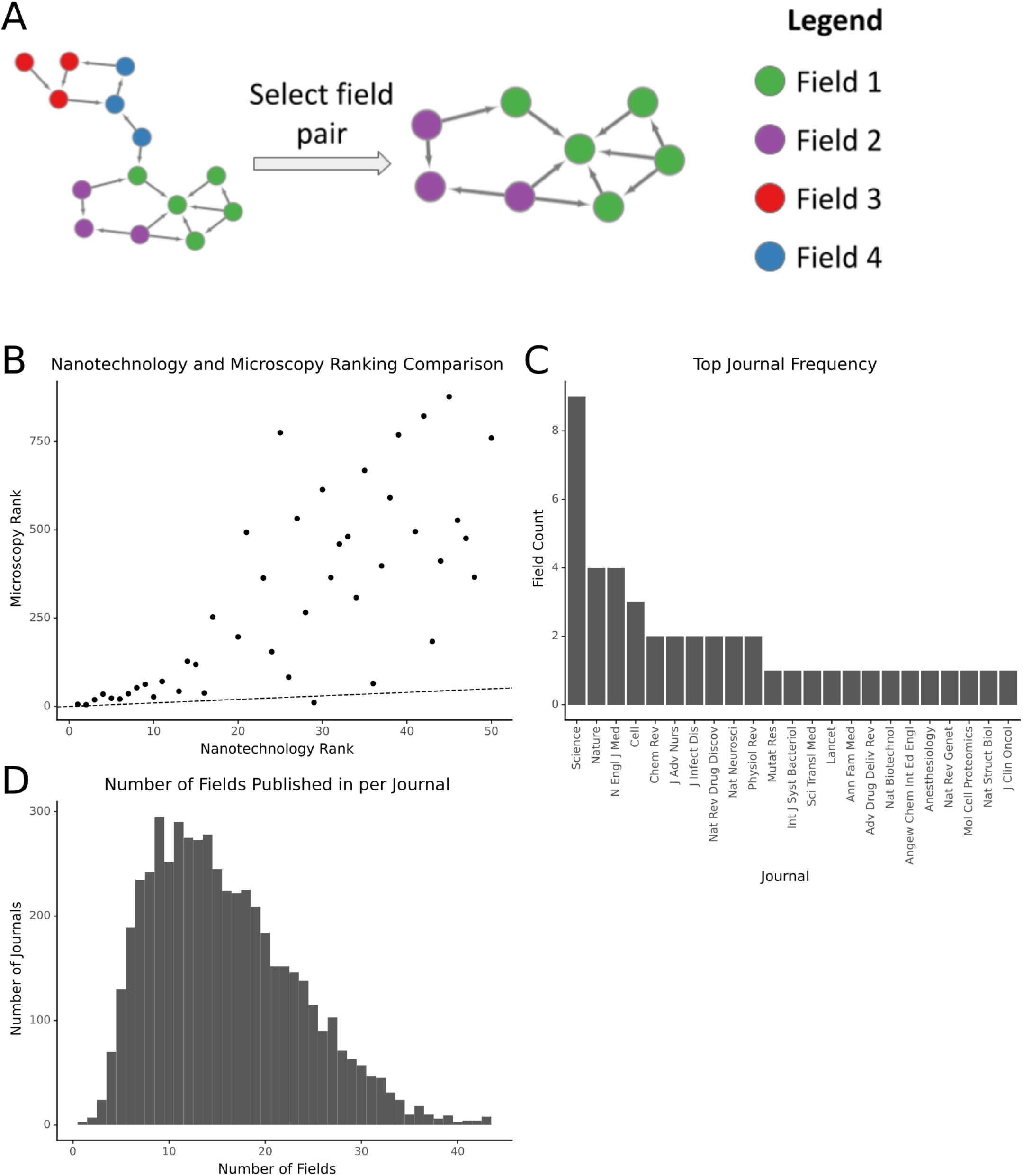
Journals’ PageRank-derived rankings differ between fields. A) A schematic showing how paired networks are derived from the full citation network. B) A comparison of the ranks of the top 50 journals by PageRank in nanotechnology and their rank in microscopy. Top-50 nanotechnology journals with no papers in microscopy have been omitted. C) The frequency with which journals in the dataset are the top journal for a field. D) The distribution of fields published per journal. The X-axis corresponds to the number of fields for which a journal has at least one paper within the field. All plots restrict the set of journals to those with at least 50 papers in the dataset.

To begin, we compared the differences in ranking between the top fifty journals in nanotechnology and their corresponding ranks in microscopy. While the ranks were correlated (r=.75) there was a great deal of variance, especially for journals outside the top 20 in nanotechnology (Fig. 1 B). We then examined the top-ranked journal in each of our 45 fields to determine whether the top-ranking journal was consistent across fields (Fig. 1 C). We found that the most commonly top-ranked journal was *Science*. This was unsurprising, given that it tends to rank highly among global journal-level metrics such as eigenfactor. However, while *Science* was the top-ranked journal in a plurality of fields, approximately 80% of fields had a different journal in that spot.

We also investigated the presence of single-topic journals in our dataset, as MeSH headings reflect a different type of aggregation than journals do [16]. Of the 5,178 journals with at least 50 articles in our dataset, the median number of fields publishing in a given journal is 15 (Fig. 1 D). In the context of MeSH, specialty journals are rare. Most journals publish manuscripts with in one-third or more of the MeSH headings in our dataset.

### Manuscript PageRanks differ between fields

We split the citation network into its component fields and calculated the PageRank for each article (Fig. 2 A). We examined the distribution of PageRanks across fields and found that they differed greatly (Fig. 2 B). We first examined whether the citation practices of fields contributed to importance differences. Investigating manuscripts that appeared in pairs of fields, we found that the distribution of importances matched the network more than that of the alternative topic area of the manuscript (Fig. 2 B, C, D).

**Figure 2:**
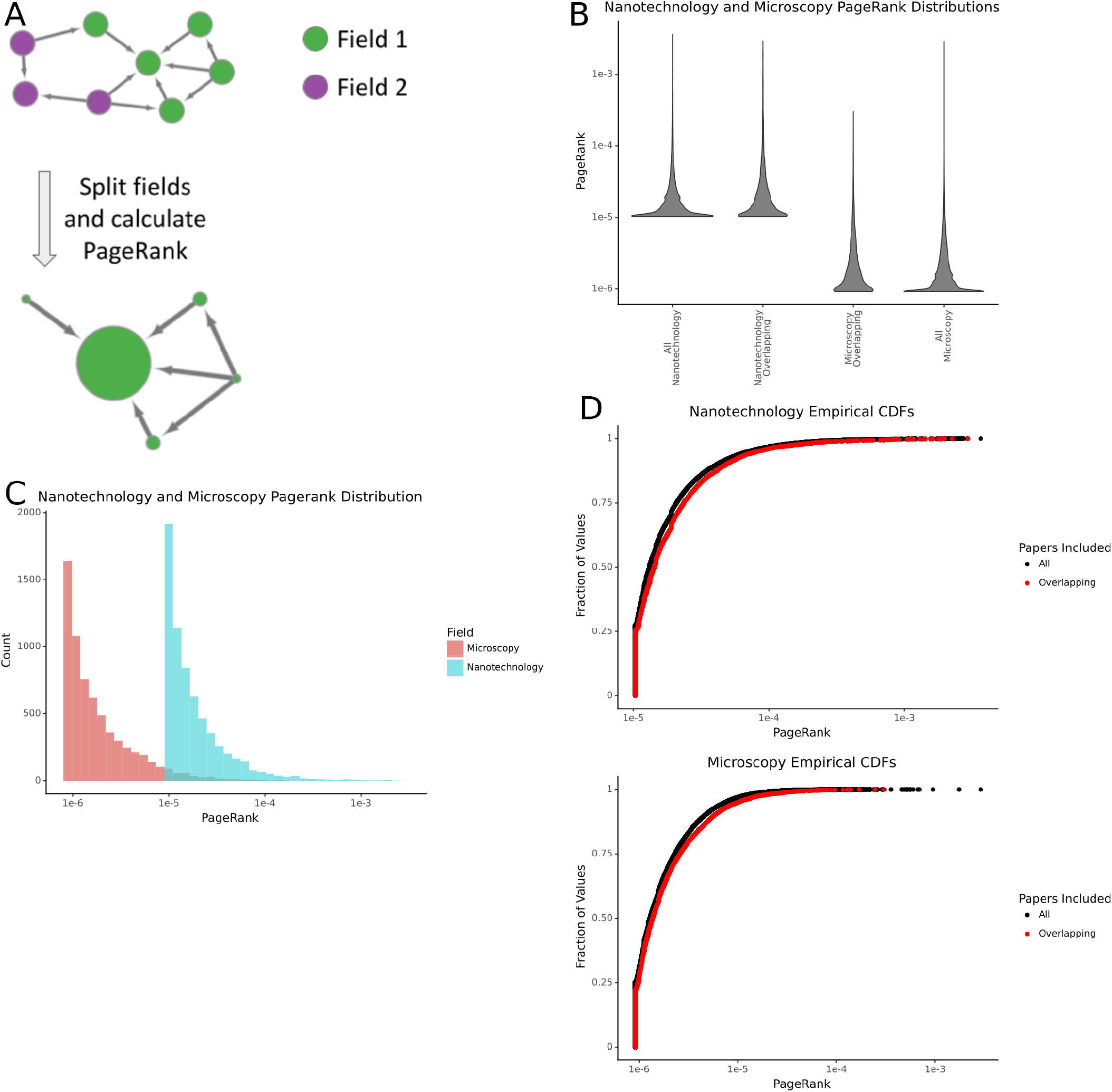
Differences in the distribution of PageRanks between fields. A) A schematic showing how field pairs are split and their PageRanks are calculated. B) The distribution of article PageRanks for nanotechnology and microscopy. The distributions marked with ‘All’ contain all the papers for the given field in the dataset, while those marked ‘overlapping’ contain only articles present in both fields. C) The empirical cumulative density functions of nanotechnology and microscopy. D) The differences in distribution of the PageRanks of articles shared by nanotechnology and microscopy. E) A density plot showing the joint distribution of PageRanks for papers overlapping in nanotechnology and microscopy.

### Fields’ differences are not solely driven by differences in citation practices

We devised a strategy to generate an empirical null for a field pair under the assumption that the field pair represented a single, homogenous field (Fig. 3 A). For each field-pair intersection, we performed a degree-distribution preserving permutation. We created 100 permuted networks for each field pair. We then split the networks into their constituent fields and calculated a percentile using the number of permuted networks with a lower PageRank for a manuscript than the true PageRank. A manuscript with a PageRank higher than all networks has a percentile of 100, and one lower than all permuted networks has a percentile of zero. We used the difference in the percentile in each field as the field-specific ainity for a given paper. This percentile score allowed us to control for the differing degree distributions between fields by comparing papers based on their expected PageRank in a random network with the same node degrees.

**Figure 3:**
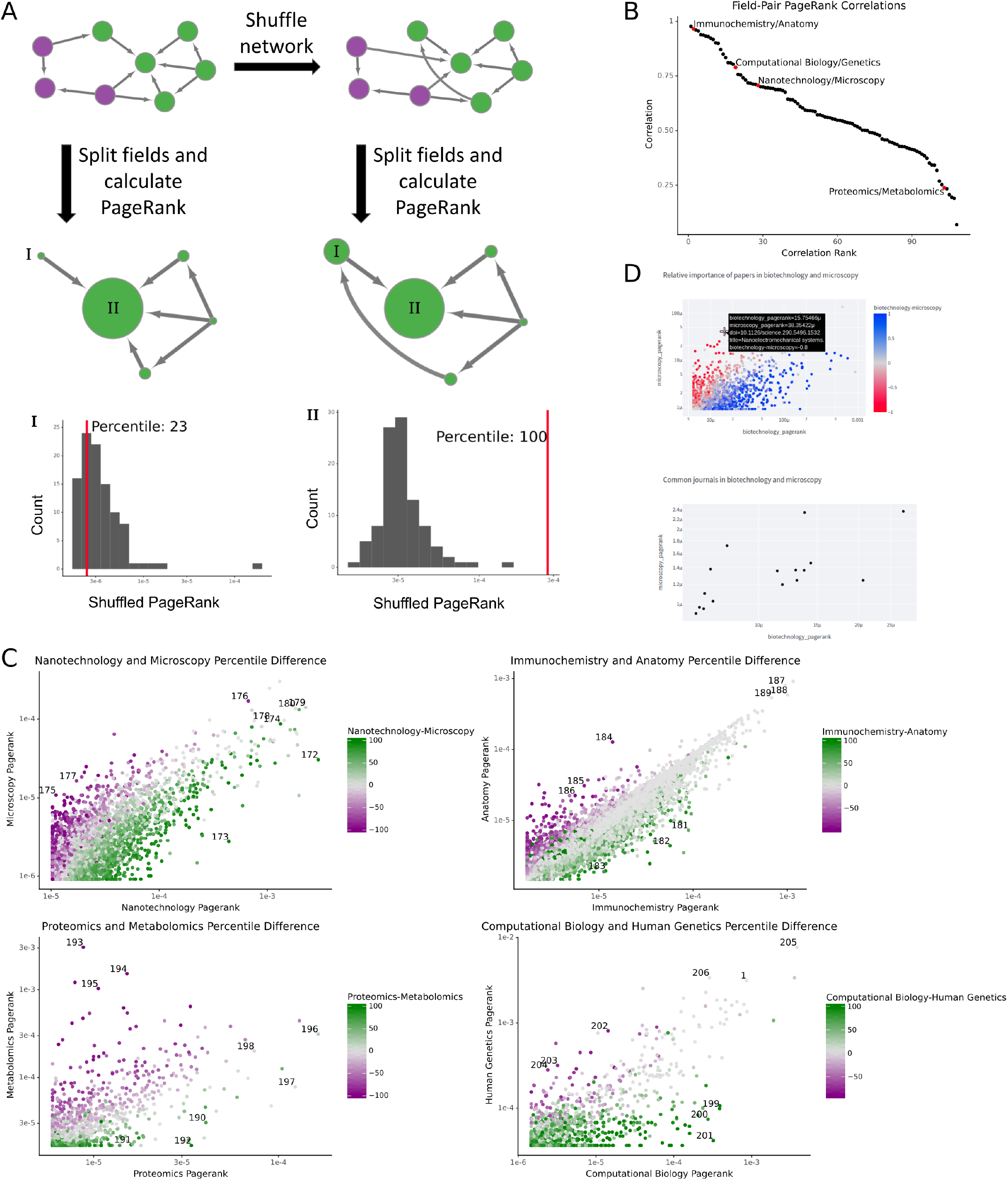
Field-specific preferences in papers. A) A schematic showing how networks are shuffled and how articles’ percentile scores are calculated. The histograms at the bottom of the figure correspond to the distribution of PageRanks for the shuffled networks, while the red lines correspond to an article’s PageRank in the true citation network. B) The Pearson correlation of PageRanks between fields. The red points are the field pairs expanded in panel C. C) The percentile scores and PageRanks for overlapping articles in various fields. Points are colored based on the difference in percentile scores in the fields e.g. “Nanotechnology-Microscopy” corresponds to the difference between the nanotechnology and microscopy percentile scores. The numbers next to points are the reference number for the article in the bibliography. D) A screenshot of the webserver showing the percentile score difference and journal median PageRank plot functionality.

We selected field pairs with varying degrees of correlation between their PageRanks (Fig. 3 B). By examining the fields’ PageRank percentiles, we found that many articles had large differences in their perception between fields (Fig. 3 C). In nanotechnology and microscopy, papers with high nanotechnology percentiles and low microscopy percentiles tended towards applications of nanotechnology, while their counterparts with high microscopy percentiles and low nanotechnology percentiles were often papers about technological developments in microscopy (Fig. 3 A, Table 1). Immunochemistry-favored papers are largely applications of immunochemical methods, while anatomy-favored articles tend to focus experiments on a single anatomical region (Fig. 3 B, Table 2). Proteomics and metabolomics tend to use similar methods, so the fields on either end are largely (though not entirely) field-specific applications of those methods (Fig. 3 C, Table 3). Manuscripts favored in computational biology were similarly applications-focused. However, those with more importance in human genetics tended towards policy papers due to its MeSH heading (H01.158.273.343.385) excluding fields like genomics, population genetics, and microbial genetics (Fig. 3 D, Table 4). In addition to papers with large differences between fields, each field pair has papers with high PageRanks and similar percentiles. While some papers may be influential in multiple fields, others have more field-specific import.

**Table 1:** Nanotechnology/microscopy papers of interest

| Nanotechnology Percentile | Microscopy Percentile | Title | Reference |
| --- | --- | --- | --- |
| 100 | 4 | A robust DNA mechanical device controlled by hybridization topology | [17] |
| 100 | 5 | Bioadhesive poly(methyl methacrylate) microdevices for controlled drug delivery | [18] |
| 99 | 2 | DNA-templated self-assembly of protein arrays and highly conductive nanowires | [19] |
| 0 | 100 | Photostable luminescent nanoparticles as biological label for cell recognition of system lupus erythematosus patients | [20] |
| 5 | 90 | WSXM: a software for scanning probe microscopy and a tool for nanotechnology | [21] |
| 0 | 77 | Measuring Distances in Supported Bilayers by Fluorescence Interference-Contrast Microscopy: Polymer Supports and SNARE Proteins | [22] |
| 100 | 99 | Toward fluorescence nanoscopy | [23] |
| 100 | 86 | In vivo imaging of quantum dots encapsulated in phospholipid micelles | [24] |
| 100 | 99 | Water-Soluble Quantum Dots for Multiphoton Fluorescence Imaging in Vivo | [25] |

**Table 2:** Immunochemistry/anatomy papers of interest

| Immunochemistry Percentile | Anatomy Percentile | Title | Reference |
| --- | --- | --- | --- |
| 100 | 45 | Immunoelectron microscopic exploration of the Golgi complex | [26] |
| 100 | 14 | Immunocytochemical and electrophoretic analyses of changes in myosin gene expression in cat posterior temporalis muscle during postnatal development | [27] |
| 98 | 5 | Electron microscopic demonstration of calcitonin in human medullary carcinoma of thyroid by the immuno gold staining method | [28] |
| 12 | 100 | Grafting genetically modified cells into the rat brain: characteristics of E. coli β-galactosidase as a reporter gene | [29] |
| 12 | 100 | Vitamin-D-dependent calcium-binding-protein and parvalbumin occur in bones and teeth | [30] |
| 3 | 100 | Mapping of brain areas containing RNA homologous to cDNAs encoding the alpha and beta subunits of the rat GABAA gamma-aminobutyrate receptor | [31] |
| 100 | 100 | Studies of the HER-2/neu Proto-Oncogene in Human Breast and Ovarian Cancer | [32] |
| 100 | 100 | Expression of c-fos Protein in Brain: Metabolic Mapping at the Cellular Level | [33] |
| 100 | 100 | Proliferating cell nuclear antigen (PCNA) immunolocalization in paraffin sections: An index of cell proliferation with evidence of deregulated expression in some neoplasms | [34] |

**Table 3:** Proteomics/metabolomics papers of interest

| Proteomics Percentile | Metabolomics Percentile | Title | Reference |
| --- | --- | --- | --- |
| 67 | 2 | Proteomics Standards Initiative: Fifteen Years of Progress and Future Work | [35] |
| 99 | 0 | Limited Environmental Serine and Glycine Confer Brain Metastasis Sensitivity to PHGDH Inhibition | [36] |
| 100 | 0 | A high-throughput processing service for retention time alignment of complex proteomics and metabolomics LC-MS data | [37] |
| 0 | 100 | MeltDB: a software platform for the analysis and integration of metabolomics experiment data | [38] |
| 0 | 98 | In silico fragmentation for computer assisted identification of metabolite mass spectra | [39] |
| 0 | 100 | The Metabonomic Signature of Celiac Disease | [40] |
| 91 | 70 | Visualization of omics data for systems biology | [41] |
| 0 | 16 | FunRich: An open access standalone functional enrichment and interaction network analysis tool | [42] |
| 0 | 5 | Proteomic and Metabolomic Characterization of COVID-19 Patient Sera | [43] |

**Table 4:**
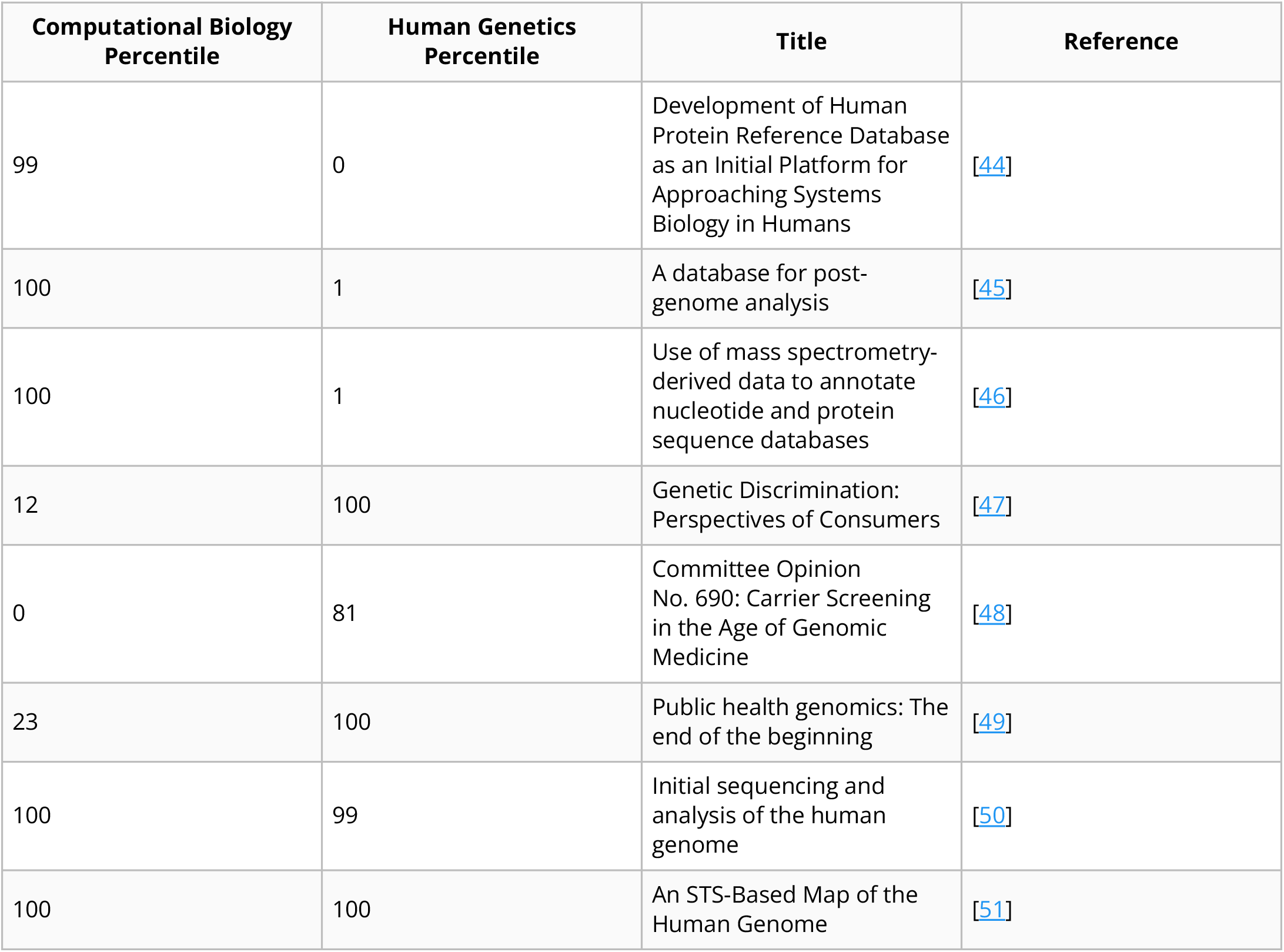
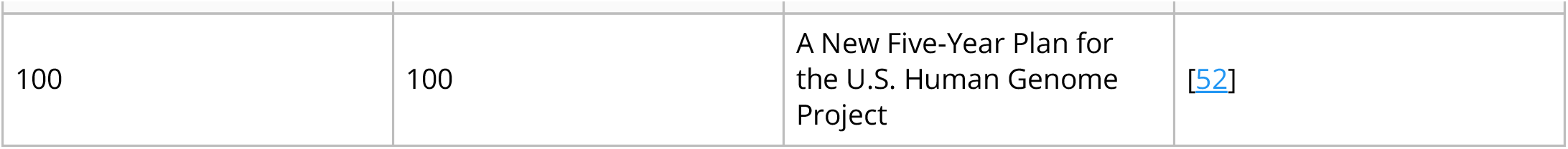
Computational biology/human genetics papers of interest

| Computational Biology Percentile | Human Genetics Percentile | Title | Reference |
| --- | --- | --- | --- |
| 99 | 0 | Development of Human Protein Reference Database as an Initial Platform for Approaching Systems Biology in Humans | [44] |
| 100 | 1 | A database for post-genome analysis | [45] |
| 100 | 1 | Use of mass spectrometry-derived data to annotate nucleotide and protein sequence databases | [46] |
| 12 | 100 | Genetic Discrimination: Perspectives of Consumers | [47] |
| 0 | 81 | Committee Opinion No. 690: Carrier Screening in the Age of Genomic Medicine | [48] |
| 23 | 100 | Public health genomics: The end of the beginning | [49] |
| 100 | 99 | Initial sequencing and analysis of the human genome | [50] |
| 100 | 100 | An STS-Based Map of the Human Genome | [51] |
| 100 | 100 | A New Five-Year Plan for the U.S. Human Genome Project | [52] |

It is impossible to describe all the field pairs and relevant differences between fields within the space of a journal article. Instead, we have developed a web server that displays the percentiles for all pairs of fields in our dataset with at least 1000 shared articles (Fig. 3 D), which can be accessed at https://www.indices.greenelab.com. We hope that the availability of the web server and the reproducibility of our code will assist other scientists in uncovering new insights from this dataset.

## Methods

### COCI

We used the March 2022 version of the COCI citation index [53] as the source of our citation data. This dataset contains around 1.3 billion citations from ∼73 million bibliographic resources.

### Selecting fields

To differentiate between scientific fields, we needed a way to map papers to fields. Fortunately, all the papers in Pubmed Central (https://www.ncbi.nlm.nih.gov/pmc/) have corresponding Medical Subject Headings (MeSH) terms. While MeSH terms are varied and numerous, the subheadings of the Natural Science Disciplines (H01) category fit our needs. However, MeSH terms are hierarchical and vary greatly in their size and specificity. To extract a balanced set of terms, we recursively traversed the tree and selected headings having at least 10000 DOIs without having multiple children that also meet the cutoff. Our resulting headings were comprised of 45 terms, from “Acoustics” to “Water Microbiology.”

### Building single heading citation networks

The COCI dataset consists of pairs of Digital Object Identifiers (DOIs). To change these pairs into a form we could run calculations on, we needed to convert them into networks. To do so, we created 45 empty networks, one for each previously selected MeSH term. We then iterated over each pair of DOIs in COCI and added them to a network if the DOIs corresponded to two journal articles written in English, both of which were tagged with the corresponding MeSH heading.

Because we were interested in the differences between fields, we also needed to build networks from pairs of MeSH headings. These networks were built via the same process, except that instead of keeping articles corresponding to a single DOI we added a citation to the network if both articles were in the pair of fields, even if the citation occurred across fields. Running this network-building process yielded 990 two-heading networks.

Sampling a graph from the degree distribution while preserving the distribution of degrees in the network was challenging. Because citation graphs are directed, it is impossible to simply swap pairs of edges and end up with a graph uniformly sampled from the space. Instead, a more sophisticated three-edge swap method must be used [54]. Because this algorithm had not been implemented yet in NetworkX [55], we implemented the code to perform shuffles and submitted our change to the library (https://github.com/networkx/networkx/pull/5663). With the shuffling code implemented, we created 100 shuffled versions of each of our combined networks to act as a background distribution against which we could compare metrics.

Once we had a collection of shuffled networks, we needed to split them into their constituent fields. To do so, we reduced the network to solely the nodes that were present in the single heading citation network and kept only citations between these nodes.

### Metrics

We used the NetworkX implementation of PageRank with default parameters to evaluate paper importance within fields. To determine the degree to which the papers’ PageRank values were higher or lower than expected, we compared the PageRank values calculated for the true citation networks to the values in the shuffled networks for each paper. We then recorded the percent of shuffled networks where the paper had a lower PageRank than the true network to derive a single number that described these values. For example, if a paper had a higher PageRank in the true network than in all the shuffled networks it received a percentile of 100. Likewise, if it had a lower PageRank in the true network than in all the shuffled networks it received a percentile of 0.

A convenient feature of the percentiles was that they were directly comparable between fields. For manuscripts represented in two fields, the difference in scores was used to estimate its variability in importance. For example, if a paper had a score of 100 in field A (indicating a higher PageRank in the field than expected given its number of citations and the network structure) and a score of 0 in field B (indicating a lower than expected PageRank), then the large difference in scores indicated the paper was more highly valued in field A than field B. If the paper had similar scores in both fields, it indicated that the paper was similarly valued in the two fields.

### Hardware/runtime

We ran the full analysis pipeline on the RMACC Summit cluster at the University of Colorado. The pipeline took about a week to run, from downloading the data to analyzing it to visualizing it. Performance in other contexts will depend heavily on details such as the number of CPU nodes available and the network speed.

### Server details

Our webserver is built by visualizing our data in Plotly (https://plotly.com/python/plotly-express/) on the Streamlit platform (https://streamlit.io/). The field pairs made available by the frontend are those with at least 1000 shared papers after filtering out papers with more than a 5% missingness level of their PageRanks after shuffling. The journals available for visualization are those with at least 25 papers for the given field pair.

## Discussion/Conclusion

We analyze hundreds of field-pair citation networks to examine the extent to which article-level importance metrics vary between fields. As previously reported, we find systematic differences in PageRanks between fields [7,56] that would warrant some form of normalization when making cross-field comparisons with global statistics. However, we also find that field-specific differences are not driven solely by differences in citation practices. Instead, the importance of individual papers appears to differ meaningfully between fields. Global rankings or efforts to normalize out field-specific effects obscure meaningful differences in manuscript importance between communities.

As with any study, this research has certain limitations. One example is our selection of MeSH terms to represent fields. We used MeSH because it is a widely-annotated set of subjects in biomedicine and thresholded MeSH term sizes to balance having enough observations to calculate appropriate statistics with having sufficient granularity to capture fields. This selection process resulted in fields at the granularity of “biophysics” and “ecology.” We also have to select a number of swaps to generate a background distribution of PageRanks for each field pair. We selected three times as many swaps as edges, where each swap modifies three edges, but certain network structures may require a different number.

We also note that there are inherent issues with the premise of ranking manuscripts’ importance. We sought to understand the extent to which such rankings were stable between fields after correcting for field-specific citation practices. We found limited stability between fields, mostly between closely-related fields, suggesting that the concept of a universal ranking of importances is difficult to justify. In the way that reducing a distribution to a Journal Impact Factor distorts assessment, attempting to use a single universal score to represent importance across fields poses similar challenges at the level of individual manucripts. Furthermore, this work’s natural progression would extend to estimating the importance of individual manuscripts to individual researchers. Thus, a holistic measure of importance would need to include a distribution of scores not only across fields but across researchers. It may ultimately be impossible to calculate a meaningful importance score. The lack of ground truth for importance is an inherent feature, not a bug, of science’s step-wide progression.

Shifting from the perspective of evaluation to discovery can reveal more appropriate uses for these types of statistics. Field-pair calculations for such metrics may help with self-directed learning of new fields. An expert in one field, e.g., computational biology, who aims to learn more about genetics may find manuscripts with high importance in genetics and low importance in computational biology to be important reads. These represent manuscripts not currently widely cited in one’s field but highly influential in a target field. Our application can reveal these manuscripts for MeSH field pairs, and our source code allows others to perform our analysis with different granularity.

## Code and Data Availability

The code to reproduce this work can be found at https://github.com/greenelab/indices. The data used for this project is publicly available and can be downloaded with the code provided above. Our work meets the bronze standard of reproducibility [57] and fulfills aspects of the silver and gold standards including deterministic operation.

## Acknowledgements

We would like to thank Jake Crawford for reviewing code that went into this project and Faisal Alquaddoomi for figuring out the web server hosting. We would also like to thank the past and present members of GreeneLab who gave feedback on this project during lab meetings. This work utilized resources from the University of Colorado Boulder Research Computing Group, which is supported by the National Science Foundation (awards ACI-1532235 and ACI-1532236).

